# Sputum Respiratory Pathogen Genomic Surveillance: A Practical Approach for Long-Read Metagenomic Sequencing

**DOI:** 10.64898/2025.12.08.692896

**Authors:** Nang Kham-Kjing, Nicole Ngo-Giang-Huong, Woottichai Khamduang

## Abstract

Severe acute respiratory infections (SARI) remain a major global health concern, particularly in resource-limited settings where comprehensive pathogen detection is challenging. Conventional diagnostics, including culture and multiplex PCR, are restricted to predefined targets and may miss emerging or uncommon pathogens. This study aimed to develop and optimize a practical long-read metagenomic next-generation sequencing (mNGS) workflow using Oxford Nanopore Technology for sputum samples from adults hospitalized with SARI. We evaluated sputum liquefaction, host nucleic acid depletion strategies, and SMART-9N-based cDNA amplification to enhance microbial and viral nucleic acid recovery. We found that dithiothreitol (DTT) treatment significantly improved nucleic acid extraction. DNase I treatment effectively reduced host background while preserving both viral and bacterial sequences, outperforming filtration-based approaches that reduced viral recovery. The optimized workflow enabled unbiased detection and strain-level resolution of respiratory pathogens, including rhinovirus C1/C42, human coronavirus HKU1, *Mycoplasma pneumoniae, Haemophilus parainfluenzae*, and *Pseudomonas aeruginosa*, with high genome coverage. This approach demonstrated robust performance for respiratory pathogen identification directly from sputum samples. The proposed workflow is scalable and suitable for clinical diagnostics and public health surveillance. Further validation in larger cohorts is warranted to assess diagnostic sensitivity, accuracy, and feasibility for routine implementation.

## Introduction

Severe acute respiratory infections (SARI) are a leading cause of hospitalization and mortality in low- and middle-income countries. SARI are caused by a wide range of respiratory pathogens, including viruses such as influenza, adenovirus, and rhinovirus, as well as bacterial species including *Klebsiella pneumoniae, Haemophilus influenzae*, and *Streptococcus pneumoniae*. Their detection rates vary substantially across populations, seasons, and geographical regions [1, 2].

Despite progress in diagnostics and immunization programs, timely and accurate identification of causative agents remains challenging. Conventional diagnostic methods such as culture, antigen detection, and multiplex polymerase chain reaction (PCR) rely on predefined pathogen panels, and as a result, they may fail to detect emerging, rare, or opportunistic pathogens, particularly in complex or polymicrobial infections [3, 4]. In resource-limited settings, these constraints are even more pronounced, underscoring the need for diagnostic approaches that are both comprehensive and adaptable. Early pathogen identification is critical not only for guiding appropriate clinical management and antimicrobial stewardship but also for enabling timely public health interventions, including outbreak detection and response.

Metagenomic next-generation sequencing (mNGS) has emerged as a transformative, hypothesis-free diagnostic approach, capable of detecting a broad spectrum of pathogens— including viruses, bacteria, fungi, and parasites—directly from clinical specimens [5]. Unlike PCR-based methods, which target specific genetic sequences, mNGS enables unbiased detection by sequencing all nucleic acids present in a sample. This allows for the identification of pathogens that may be unculturable, unexpected, or absent from conventional diagnostic panels [6, 7]. mNGS is particularly well-suited for complex syndromes such as respiratory infections, which often involve multiple pathogen types coexisting within the same sample [8].

To date, the majority of mNGS applications are short-read platforms such as Illumina. While highly accurate, these platforms are limited in their ability to resolve repetitive or homologous regions, detect structural variants, or generate complete genome assemblies [9]. In contrast, long-read sequencing technologies, particularly those based on Oxford Nanopore Technologies (ONT), offer advantages such as real-time sequencing, extended read lengths, and near-complete genome reconstruction [9, 10]. Long-read technologies enable in-depth phylogenetic analysis, strain-level identification, and improved surveillance of antimicrobial resistance genes [11, 12].

Nevertheless, the application of long-read metagenomic sequencing for the identification of respiratory pathogen detection in sputum remains limited. Sputum is the specimen of choice for diagnosing lower respiratory tract infections in adults. However, it poses significant technical barriers to sequencing workflows. Indeed, sputum is characterized by high viscosity due to the mucin content, the abundance of host-derived nucleic acids, and the presence of PCR inhibitors, which complicate nucleic acid extraction, depletion of the host genome, and decreased efficiency of downstream sequencing. While ONT-based mNGS has been successfully applied to nasopharyngeal and bronchoalveolar samples [13, 14], there is a critical lack of optimized protocols specifically tailored for sputum specimens.

In this study, we aimed to develop and optimize a long-read mNGS workflow for sputum samples collected from adults hospitalized with SARI. Specifically, we evaluated multiple sputum processing and host genome depletion strategies to enhance microbial and viral nucleic acid yield while minimizing host background. Additionally, we refined library preparation and cDNA amplification protocols to support high-quality, untargeted long-read sequencing. The resulting workflow provides a practical and scalable approach for pathogen surveillance and genomic epidemiology in respiratory diseases, with potential applications in both clinical diagnostics and public health preparedness.

## Results

### Evaluation of sputum pretreatment for mNGS

To determine the impact of pretreatment methods on nucleic acid recovery and downstream pathogen detection, three sputum processing protocols were compared: untreated control, treatment with Zybio pretreatment solution, and 0.1% dithiothreitol (DTT). The HRV-positive sample pretreated with DTT yielded the lowest cycle threshold (Ct) value for human rhinovirus (Ct = 17.1) as compared to the untreated control (Ct = 21.8) and Zybio pretreatment (Ct = 31.6). Furthermore, the internal control (IC) Ct values in DTT-treated samples were consistently lower by 4–6 cycles compared to untreated samples and 0–3 cycles compared to Zybio-treated samples. These results indicate that DTT pretreatment of sputum enhanced extraction efficiency and reduced PCR inhibition (Fig. 1).

**Figure 1.**
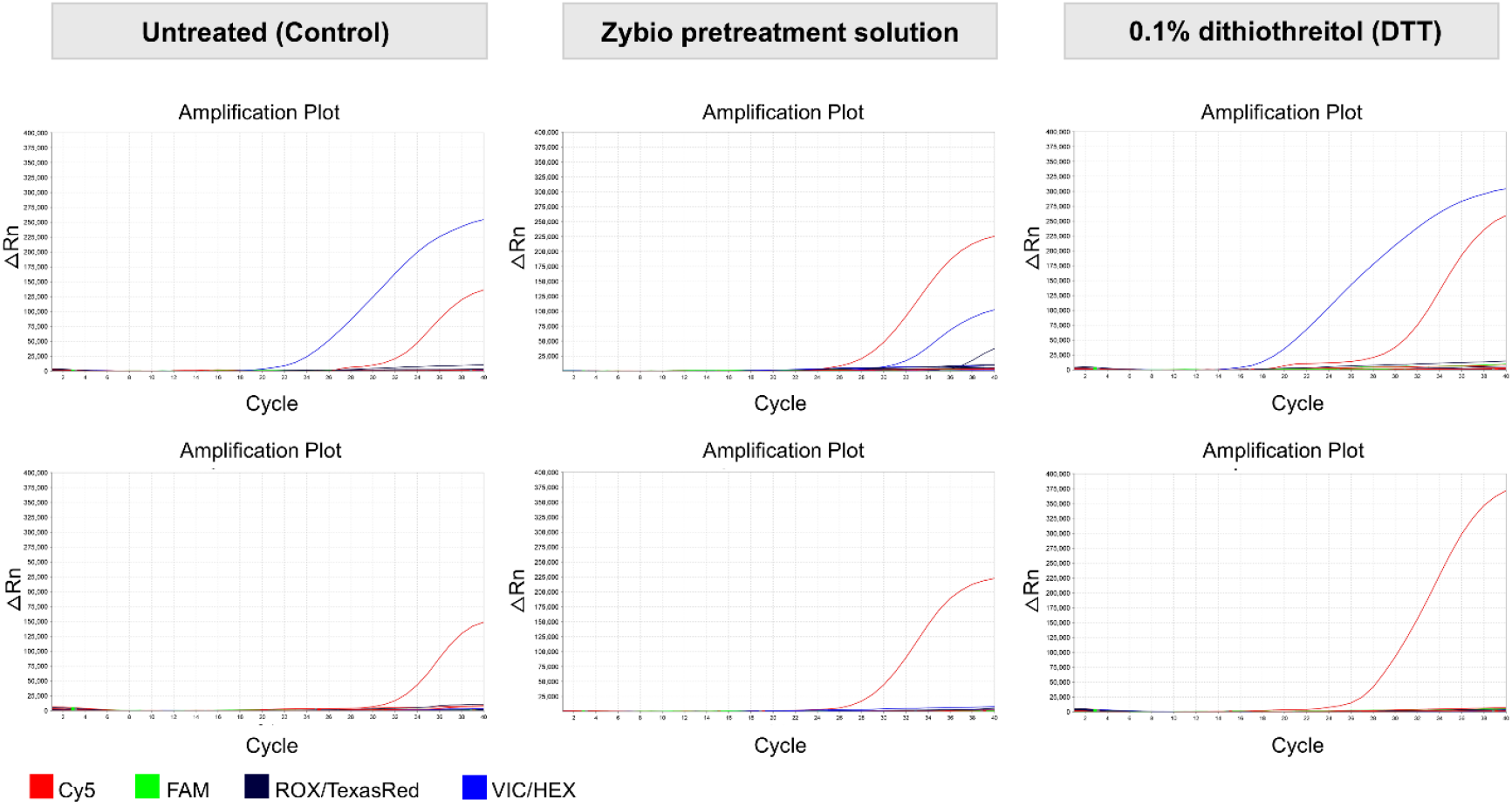
Comparison of RNA extracts after no, Zybio, and DTT pretreatment of sputum using multiplex real-time RT-PCR. Blue line: HRV-positive signal; Red line: internal control.

### Enhancing microbial enrichment by host genome or DNA depletion

To reduce host DNA background and enhance microbial signal, three host DNA depletion strategies were evaluated alongside an untreated control: DNase I treatment alone, combined filtration and DNase I, and adaptive sampling. Untreated specimens yielded nearly 100% host-derived reads, indicating minimal microbial content (Fig. 2a and Supplementary Fig. S1-S2). Both treatment with DNase I alone and the combined approach substantially reduced host reads and most effectively enriched bacterial sequences (Fig. 2b, 2c, Supplementary Fig. S1-S5). However, enterovirus and human coronavirus HKU1 (each constituting ∼13% of classified reads) were only detected in samples processed with the DNase I-only strategy (Supplementary Fig. S4). Moreover, treatment with DNase I alone markedly increased the proportion of viral reads compared with other methods, as shown in Figure 2. Adaptive sampling demonstrated potential for real-time host read rejection and microbial enrichment (Fig. 2d), but its implementation is limited by the need for powerful computational hardware. A detailed comparison of sequencing metrics for host depletion efficiency across the six samples is provided in Supplementary Table S1.

**Figure 2.**
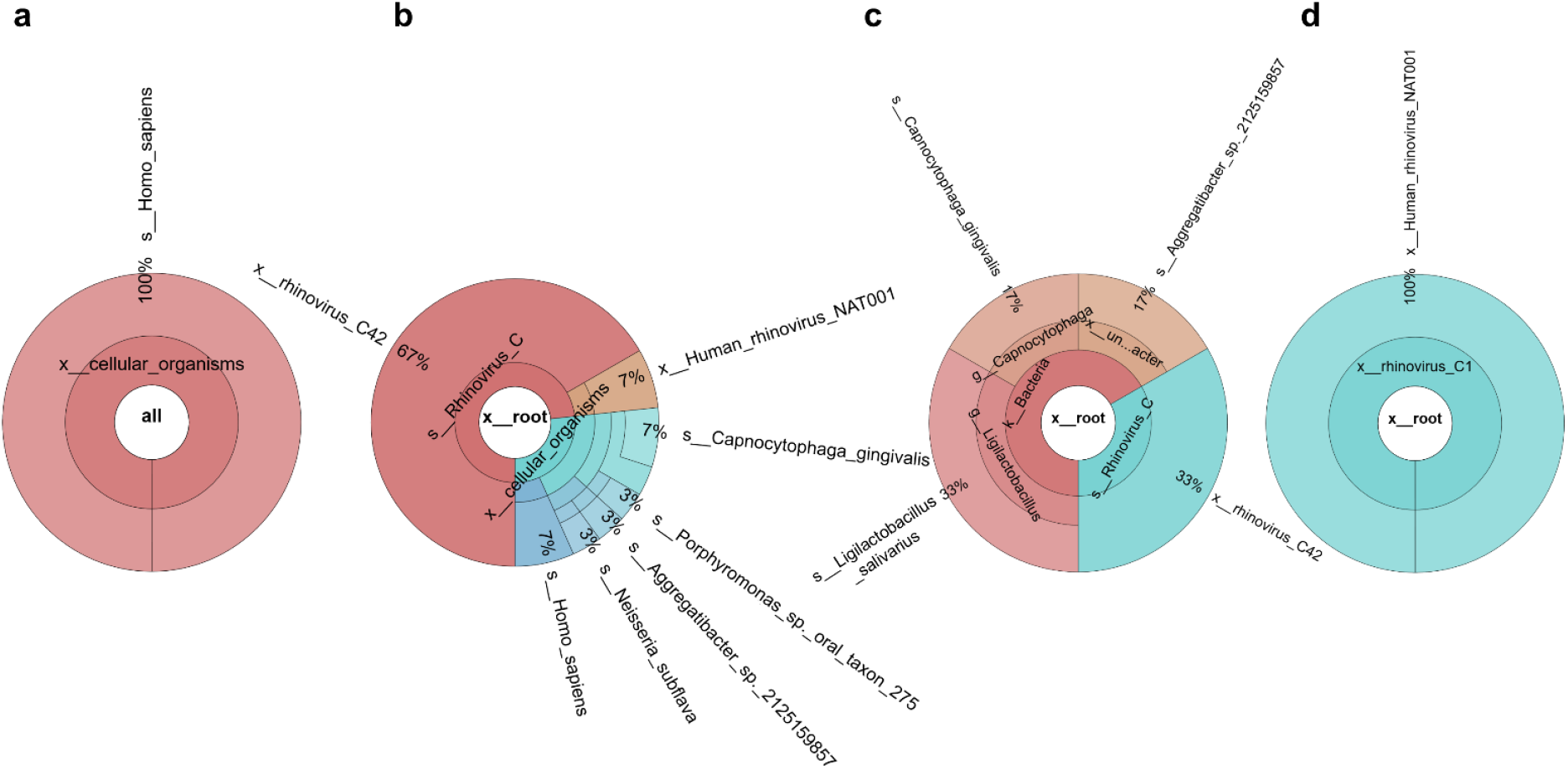
Comparison of host DNA depletion strategies: (a) untreated; (b) DNase I only; (c) filtration + DNase I; (d) adaptive sampling.

### Optimal sputum metagenomic sequencing workflow

Based on experimental comparisons, an optimized workflow was established (Fig. 3). This integrated approach utilized 0.1% DTT for efficient sputum liquefaction, DNase I for selective host DNA depletion, SMART-9N for full-length cDNA amplification, and ONT sequencing for long-read analysis. The optimized workflow enabled taxonomic identification of respiratory pathogens down to the species level, and in all 6 cases, resolution extended to the strain or genotype level. Detected pathogens included rhinovirus C, human coronavirus HKU1, *Mycoplasmoides pneumoniae, Haemophilus parainfluenzae, Pseudomonas aeruginosa, Pseudomonas putida*, and *Metamycoplasma salivarium*. Strain-level classifications included rhinovirus C1, rhinovirus C42, and human rhinovirus NAT001.

**Figure 3.**
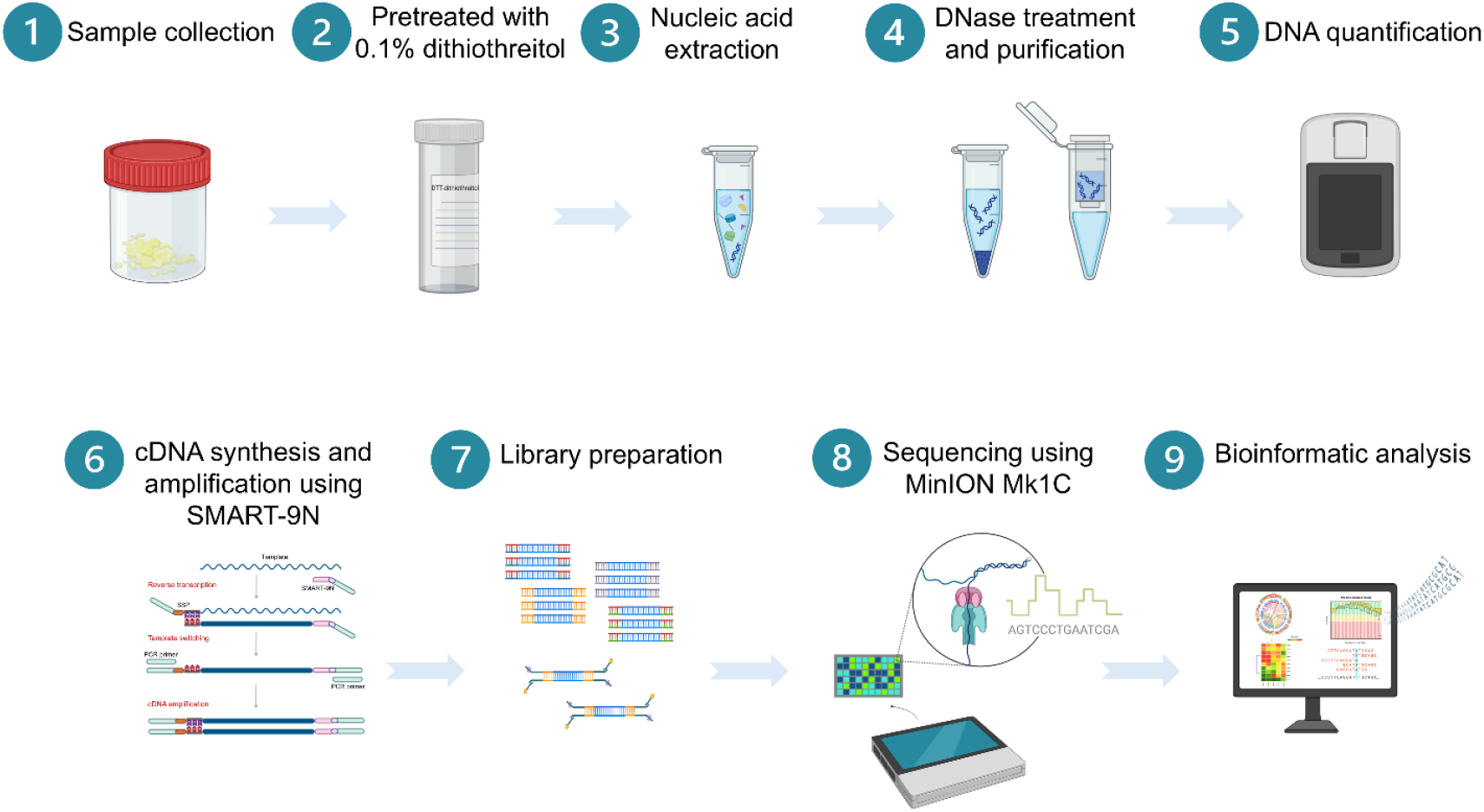
Overview of the optimized long-read sequencing workflow for detection of respiratory pathogen in sputum.

### Genome coverage

Genome coverage metrics were assessed to evaluate sequencing depth and completeness across representative respiratory pathogens. Human coronavirus HKU1 displayed uniform genome-wide coverage, with average sequencing depth exceeding 40×. Human rhinovirus demonstrated even greater depth, with multiple regions surpassing 400×. Bacterial pathogens also showed high coverage, with *Mycoplasma pneumoniae* reaching over 15,000× depth in some regions (Fig. 4).

**Figure 4.**
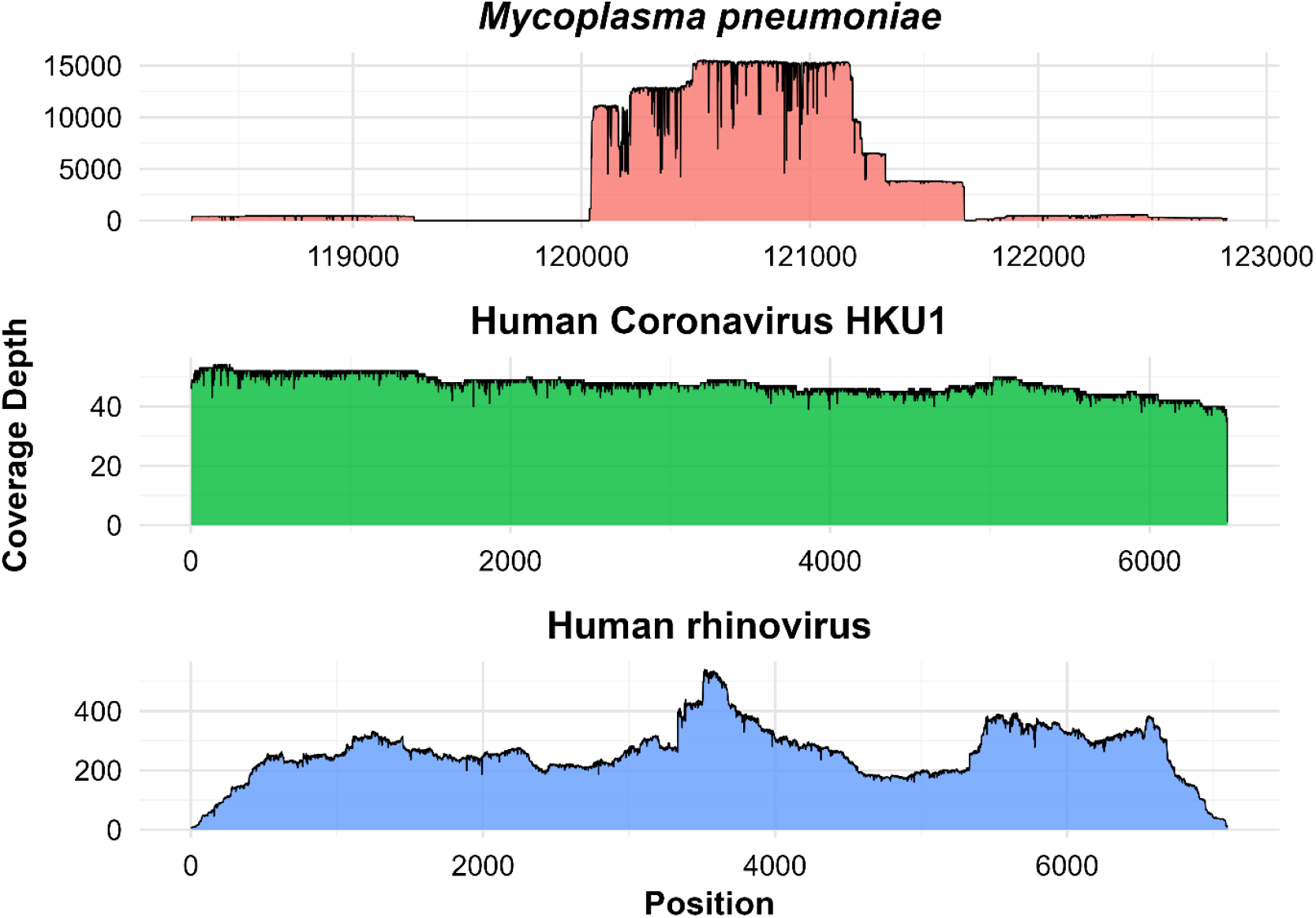
Genome coverage plots showing key respiratory pathogens identified in SARI sputum samples. Coverage depth (y-axis) is plotted against genome position (x-axis); the scales of the axes vary by plot.

## Discussion

This study presents a systematically optimized long-read metagenomic sequencing (mNGS) workflow specifically tailored for sputum samples from patients hospitalized with SARI. The integrated protocol combines 0.1% dithiothreitol (DTT) for sputum liquefaction, DNase I treatment for host nucleic acid depletion, SMART-9N amplification for cDNA synthesis, and Oxford Nanopore Technologies (ONT) sequencing, yielding improved nucleic acid recovery and enabling high-resolution pathogen detection at the species and strain/genotype levels (e.g., rhinovirus C1/C42, *Haemophilus parainfluenzae*, and *Pseudomonas aeruginosa*).

Among the evaluated pretreatment strategies, DTT consistently yielded lower Ct values in multiplex real-time RT-PCR compared to untreated and Zybio-treated samples, indicating improved microbial nucleic acid release and PCR performance. These findings are consistent with previous reports showing that DTT homogenizes viscous sputum by reducing mucin disulfide bonds, thereby improving DNA extraction yields and reducing variability across replicates [15, 16]. Additionally, studies comparing DTT with proteinase K and normal saline have demonstrated that DTT is superior in enhancing PCR detection sensitivity for key respiratory pathogens in sputum samples [17, 18]. Given its simplicity, low cost, and compatibility with downstream applications, DTT represents a highly practical pretreatment option for laboratories working with lower respiratory tract specimens.

In the host depletion step, a trade-off was observed between bacterial enrichment and viral detection. The combined filtration and DNase I approach significantly reduced host-derived reads and enhanced bacterial signal. However, it compromised viral detection, likely due to the loss of viral particles during filtration. In contrast, DNase I treatment alone effectively reduced host background while preserving both bacterial and viral nucleic acids, enabling the detection of human coronavirus HKU1 and enterovirus, which were undetectable under the combined protocol. These results are consistent with previous reports showing that nuclease-based depletion methods enrich viral-to-host read ratios without significantly degrading viral content [19]. These results highlight the need for tailoring host depletion strategies according to the surveillance objective— bacterial vs. viral focus. Importantly, our optimized workflow enabled the recovery of near-complete viral genomes and provided species- and strain-level taxonomic resolution, a level of resolution essential for outbreak detection, molecular epidemiology, and tracking of viral evolution.

Beyond our findings, recent studies have reinforced the importance of host DNA depletion strategies in improving metagenomic sequencing of respiratory samples. For instance, Kim *et al*. evaluated 5 different host DNA depletion methods (lyPMA, Benzonase, HostZERO, MolYSIS, and QIAamp) in frozen respiratory specimens, showing that host depletion markedly increased microbial detection and species richness [20]. Moreover, Hasan *et al*. demonstrated that saponin lysis followed by DNase treatment achieves robust host DNA removal and enriches pathogens in nasopharyngeal aspirates and cerebrospinal fluid [21].

Nevertheless, this study has limitations. The evaluation was conducted on a limited number of samples. Future studies should incorporate larger, diverse cohorts to validate sensitivity, specificity, and reproducibility. Furthermore, comparative studies evaluating cost-effectiveness and turnaround time between our workflow and conventional diagnostic methods will be essential for assessing feasibility in clinical and public health surveillance contexts.

## Material and Methods

### Sample collection

Sputum specimens were collected from hospitalized adult patients diagnosed with severe acute respiratory infection (SARI) at the regional hospital in Chiang Mai, Thailand. SARI criteria were defined as acute onset of respiratory symptoms within the past 10 days, measured or reported fever ≥38°C, presence of cough, and clinical requirement for hospitalization. Collected samples were transported on ice to the central laboratory and stored at –70°C until further processing.

### Sample pretreatment

Two sputum samples were selected for the mNGS optimization experiments based on their previously tested multiplex PCR results using the FTD™ Respiratory Pathogens 21 multiplex real-time RT-PCR assay (Fast Track Diagnostics, Siemens Healthineers, Esch-sur-Alzette, Luxembourg). One sample had been confirmed positive for human rhinovirus (HRV), while the other had tested negative for all respiratory pathogens included in the panel. Each of the two samples was subsequently aliquoted into three tubes and subjected to three different pretreatment conditions:

i. Untreated control: Samples were centrifuged at 10,000 × g for 3 minutes, and the supernatant was retained.
ii. Zybio pretreatment: 200 µL of sputum was mixed with 100 µL of pretreatment buffer from the Zybio Nucleic Acid Extraction Kit BF-B-32 (Zybio, Chongqing, China), incubated at 90°C for 10 minutes, centrifuged, and the supernatant was collected.
iii. Dithiothreitol (DTT) treatment: An equal volume of 0.1% DTT (Thermo Fisher Scientific, MA, USA) was added to the sputum and incubated at room temperature until complete liquefaction. The mixture was centrifuged at 10,000 × g for 3 minutes, and the supernatant was used for nucleic acid extraction.

### Nucleic acid extraction and evaluation of pretreatment effects

Total nucleic acid was extracted from the pretreated sputum samples using the QIAamp® Viral RNA Mini Kit (Qiagen GmbH, Hilden, Germany), according to the manufacturer’s instructions. All extractions were performed in parallel to maintain consistency. To assess the impact of pretreatment conditions on pathogen detection, nucleic acid extracts were evaluated using the FTD^™^ Respiratory Pathogens 21 multiplex real-time RT-PCR assay (Fast Track Diagnostics, Siemens Healthineers, Esch-sur-Alzette, Luxembourg). Cycle threshold (Ct) values were compared for the optimal pretreatment conditions.

### Host genome removal

To reduce host genome background, two host depletion strategies were tested:

i. Untreated control: Extracted nucleic acids were subjected to cDNA synthesis and amplified directly without any treatment.
ii. DNase I treatment only: Extracted nucleic acids were treated with RNase-Free DNase Set (Qiagen GmbH, Hilden, Germany) and purified using the RNA Clean & Concentrator™-25 Kit (Zymo Research, CA, USA).
iii. Filtration followed by DNase I treatment: Pretreated sputum was first filtered through a 0.45 µm syringe filter (Minisart^®^ NML, Sartorius Stedim Biotech GmbH, Göttingen, Germany) to remove eukaryotic cells and debris. Filtrates were subjected to nucleic acid extraction, followed by DNase I treatment as above.
iv. Adaptive sampling: DNase I treated extracts were used for sequencing library preparation, and real-time selective sequencing (adaptive sampling) with negative selection on the human reference genome (GRCh38) was enabled during sequencing.

### cDNA synthesis and amplification using SMART-9N technology

cDNA synthesis and amplification were performed using SMART-9N (**S**witching **M**echanism **a**t the 5′ end of **R**NA **T**emplate with random nonamer primers) technology, adapted from a previously described protocol [22]. This method utilizes a specialized reverse transcriptase capable of template-switching via a strand-switching primer (SSP) containing a 3′ rGrGrG motif, enabling full-length cDNA synthesis with universal priming sites (Fig. 5).

**Figure 5.**
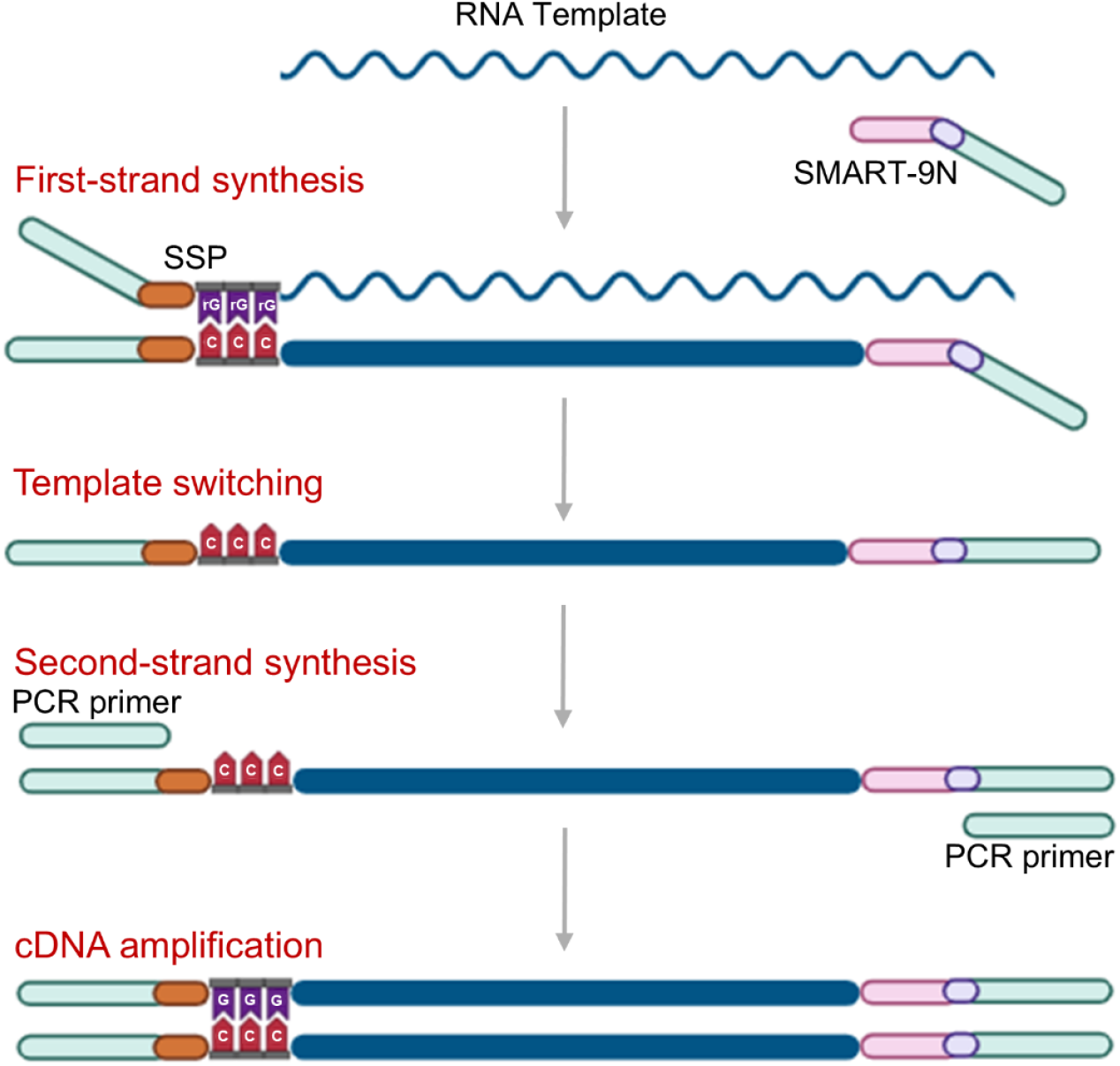
Overview of the SMART-9N cDNA synthesis and amplification workflow.

For cDNA synthesis, 1 µL of 2 µM 9N primer (AAGCAGTGGTATCAACGCAGAG TACNNNNNNNNN), 1 µL of 10 mM dNTP mix, and 10 µL of purified RNA were combined, heated to 65°C for 5 minutes, and chilled on ice. The first-strand synthesis included 4 µL of 5× SuperScript™ IV buffer, 1 µL each of 0.1 M DTT, RNase inhibitor, SuperScript™ IV reverse transcriptase (Thermo Fisher Scientific, MA, USA), and 2 µM SSP (GCTAATCATTGCAAGCAGTGGTATCAACGCAGAGTACATrGrGrG), along with 12 µL of annealed RNA. The mixture was incubated at 42°C for 90 minutes, followed by 70°C for 10 minutes.

PCR amplification was performed using 2.5 µL of cDNA in a total 25 µL reaction containing 5 µL of Q5® reaction buffer, 0.5 µL of 10 µM dNTPs, 1 µL of 20 µM PCR primer (AAGCAGTGGTATCAACGCAGAGT), and 0.25 µL of Q5® High-Fidelity DNA Polymerase (New England Biolabs, MA, USA). Cycling conditions were as follows: initial denaturation at 98°C for 45 seconds; 30 cycles of 98°C for 15 seconds, 62°C for 15 seconds, and 65°C for 5 minutes; and final extension at 65°C for 10 minutes. Amplicons were purified using VAHTS DNA Clean Beads (Vazyme, Nanjing, China) at a 1:1 ratio. DNA quantification was performed using the DeNovix dsDNA Broad Range Assay Kit and measured on a QFX Fluorometer (DeNovix Inc., DE, USA).

### Metagenomic next-generation sequencing

Sequencing libraries were prepared using 50 ng of purified amplicons with the Native Barcoding Kit 24 V14 (ONT-SQK-NBD114.24) following Oxford Nanopore Technologies protocols. Libraries were quantified, pooled, and loaded onto the MinION Flow Cell R10.4.1 version (FLO-MIN114 flow cells). Sequencing was conducted on a MinION Mk1C platform (Oxford Nanopore Technologies, Oxford, UK) for 72 hours.

### Bioinformatics analysis

Raw sequencing data (pod5 files) generated by the MinION Mk1C platform were basecalled using Dorado with a high-accuracy model with the MinKNOW v24.11.10 software and retaining reads with a minimum average quality score of Q9. Demultiplexing and adapter trimming were also performed using the MinKNOW software. Quality control analysis was performed using NanoComp v1.23.1, and the detailed sequencing metrics were visualized using NanoPlot v1.42.0 [23]. The SMART-9N primer overhangs were trimmed using Cutadapt v5.0 [24] with a maximum error rate of 10% and a minimum overlap of 25 base pairs (bp) without indels, followed by quality filtering using fastp v0.24 [25], retaining reads with a minimum length of 200 bp. Sequencing reads were aligned to the human reference genome (GRCh38) using Minimap2 v2.28-r1209 [26] with Oxford Nanopore–specific settings (-x map-ont) and a quality occupancy fraction of 0.01 to remove low-confidence host mappings. Host-derived reads were filtered out, and the unmapped reads were extracted for downstream analysis. De novo assembly of these host-depleted reads was performed using Flye v2.9.5-b1801 [27] with the nano-hq option in meta mode and two iterations. The resulting contigs were polished with Medaka v2.0.1 [28] to improve base-level accuracy and generate consensus sequences. Taxonomic classification was performed with Kraken2 v2.1.3 [29] with the PlusPF database, which combines the standard Kraken2 database with RefSeq protozoan and fungal genomes. Both classified and unclassified reads were retained for downstream analyses. Relative abundances were calculated from classified read counts and visualized using KronaTools v2.8.1 [30]. Read mapping and genome coverage analyses were performed using Minimap2. Coverage depth plots were generated in RStudio v4.4.2 (RStudio, MA, USA) using the ggplot2 package.

### Ethical approval

This study was approved by the Ethics Committee of the Faculty of Associated Medical Sciences, Chiang Mai University (Approval No. AMSEC-66EM-011), and the regional hospital ethics board (Approval No. 112/66). All specimens were anonymized, and residual samples from routine diagnostics were used under a waiver of informed consent, in accordance with ethical guidelines.

## Conclusions

Severe acute respiratory infections remain a major public health concern, particularly in settings with limited diagnostic infrastructure. This study addresses critical gaps in pathogen detection from sputum by establishing a robust and reproducible long-read metagenomic workflow. The integration of DTT-based sample liquefaction, DNase I host depletion, and SMART-9N amplification, combined with ONT sequencing, enabled high-resolution identification of respiratory pathogens, including strain-level classification of viruses and bacteria. This optimized protocol offers a scalable solution for unbiased pathogen surveillance, enhancing diagnostic accuracy and supporting outbreak preparedness. Further studies involving larger populations are warranted to validate the performance and evaluate its integration into clinical diagnostic workflows and national surveillance programs.

## Acknowledgements

N.K.K. is a candidate in the PhD Degree Program in Biomedical Sciences, Faculty of Associated Medical Sciences (AMS), Chiang Mai University (CMU), under the CMU Presidential Scholarship. We gratefully acknowledge Nakornping Hospital for providing clinical samples and the Research Institute for Health Sciences and the LUCENT International Collaboration, AMS, CMU for their laboratory support and facilities.

## Author contributions

Conceptualization: W.K.; Methodology: N.K.K.; Software: N.K.K.; Sequence data analysis: N.K.K.; Validation: N.N. and W.K.; Investigation: all; Resources: N.N. and W.K.; Visualization: N.K.K.; Supervision: N.N. and W.K.; Funding Acquisition: W.K.; Original Draft Preparation: N.K.K.; Writing – Review and Editing: all.

