## Supplementary data for "Sputum Respiratory Pathogen Genomic Surveillance: A Practical Approach for Long-Read Metagenomic Sequencing"

| **Sample** | **Methods** | **% Host read** | **% Microbial read** | **Unique microbial species detected** | |
| --- | --- | --- | --- | --- | --- |
|  |  |  |  | **Bacteria (n%)** | **Viruses (n%)** |
| S1 | Control | 100% | - | - | - |
|  | Dnase I | 7% | 93% | 4 (20%) | 1 (73%) |
|  | Filtration + Dnase I | - | 100% | 3 (67%) | 1 (33%) |
|  | Adaptive sampling | - | 100% | - | 1 (100%) |
| S2 | Control | 100% | - | - | - |
|  | Dnase I | - | 100% | 1 (25%) | 1 (75%) |
|  | Filtration + Dnase I | NA | NA | NA | NA |
|  | Adaptive sampling | - | 100% | 1 (100%) | - |
| S3 | Control | 100% | - | - | - |
|  | Dnase I | - | 100% | 3 (100%) | - |
|  | Filtration + Dnase I | - | 100% | 1 (100%) | - |
|  | Adaptive sampling | NA | NA | NA | NA |
| S4 | Control | 1% | 95% | 14 (95%) | - |
|  | Dnase I | - | 100% | - | 1 (100%) |
|  | Filtration + Dnase I | - | 100% | - | 1 (100%) |
|  | Adaptive sampling | - | 100% | 1 (100%) | - |
| S5 | Control | - | 100% | 5 (100%) | - |
|  | Dnase I | 13% | 87% | 4 (62%) | 2 (25%) |
|  | Filtration + Dnase I | - | 100% | 5 (100%) | - |
|  | Adaptive sampling | NA | NA | NA | NA |
| S6 | Control | 50% | 50% | 1 (50%) | - |
|  | Dnase I | - | 100% | 2 (100%) | - |
|  | Filtration + Dnase I | 25% | 75% | 3 (75%) | - |
|  | Adaptive sampling | - | 100% | 1 (100%) | - |

**Supplementary Table 1. Summary of sequencing metrics for host depletion optimization.**

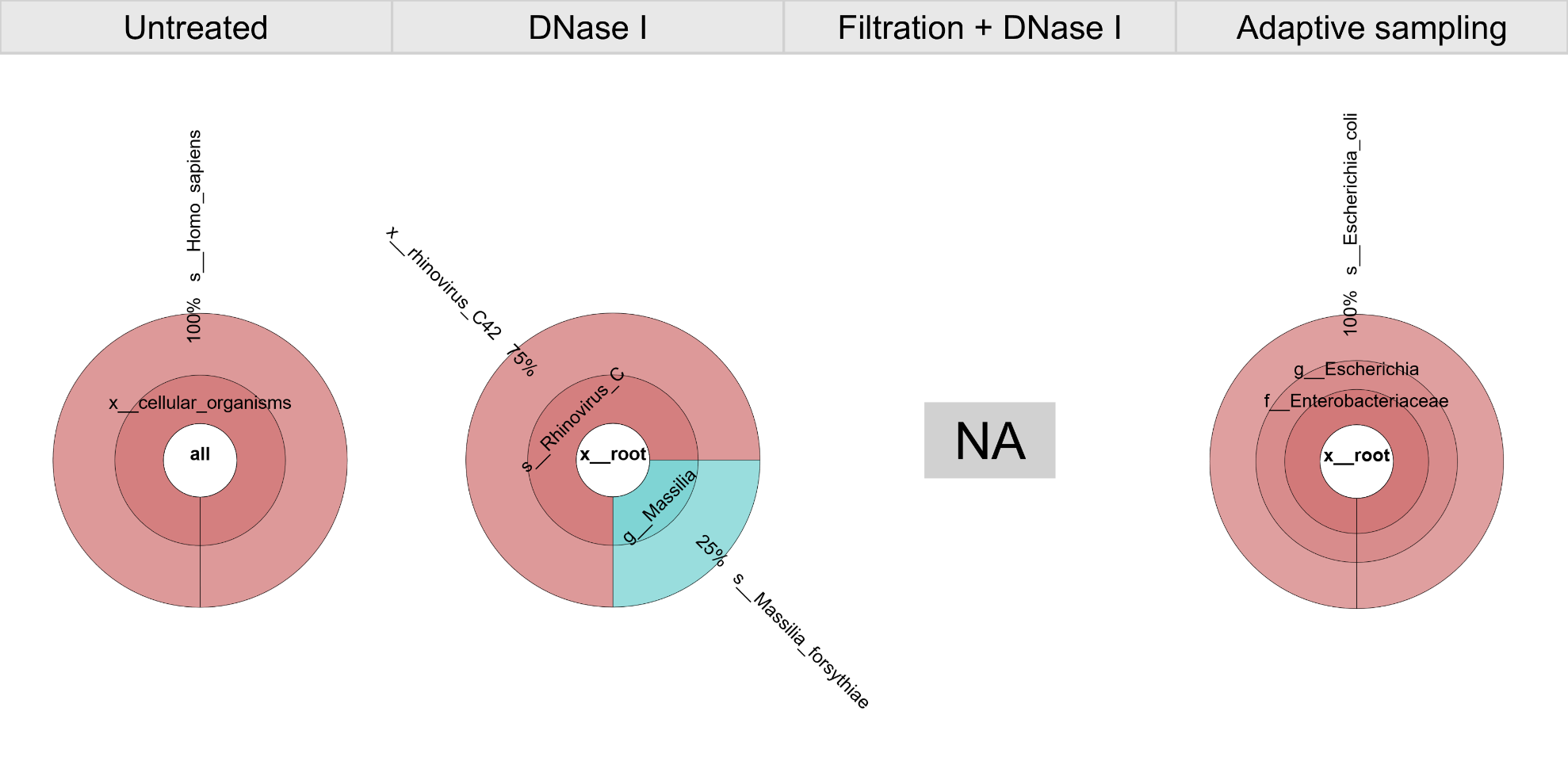

**Supplementary Figure S1.** Taxonomic composition of sample S2 following four host depletion strategies visualized using Krona plots. NA: not available.

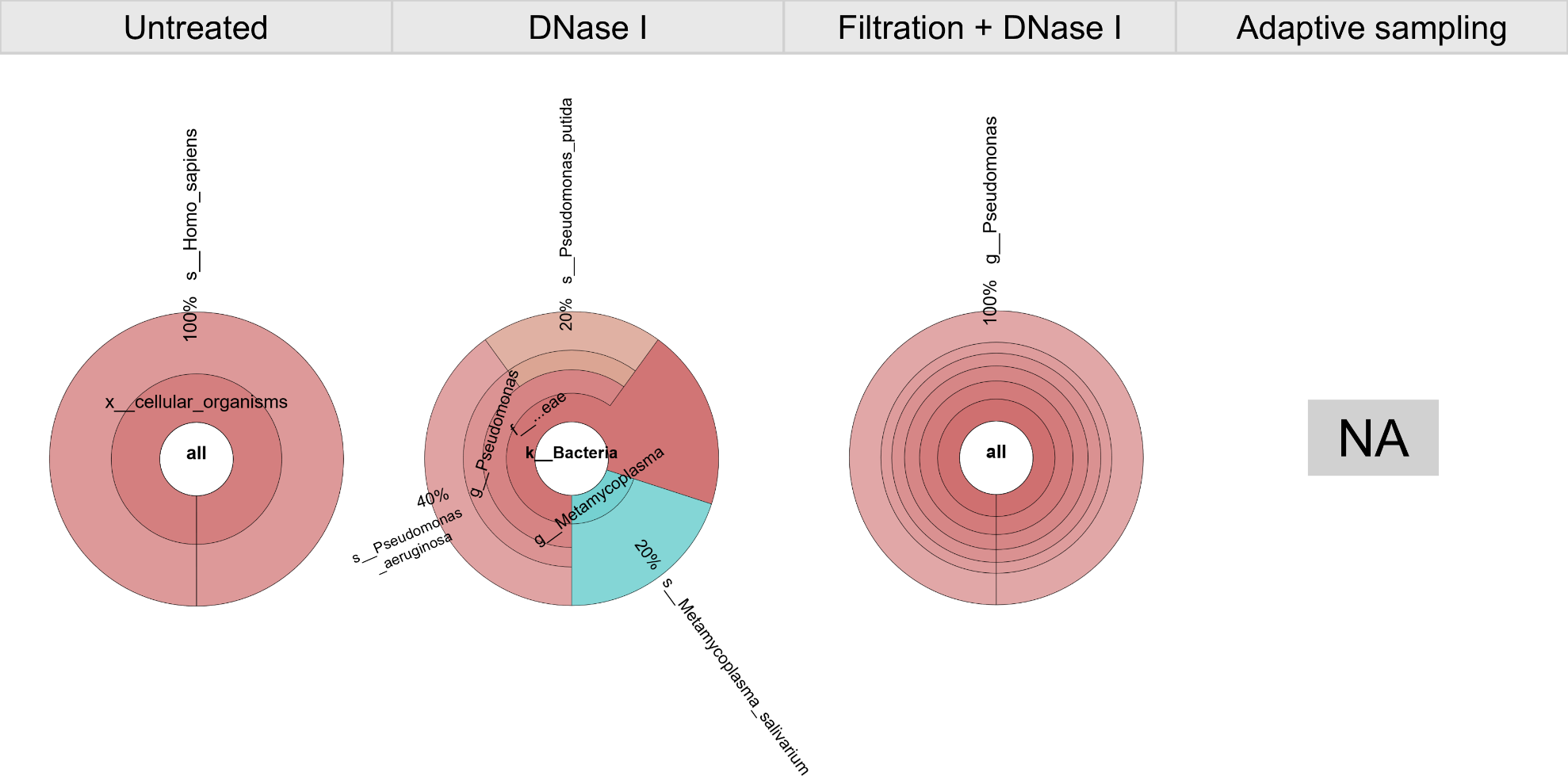

**Supplementary Figure S2.** Taxonomic composition of sample S3 following four host depletion strategies visualized using Krona plots. NA: not available.

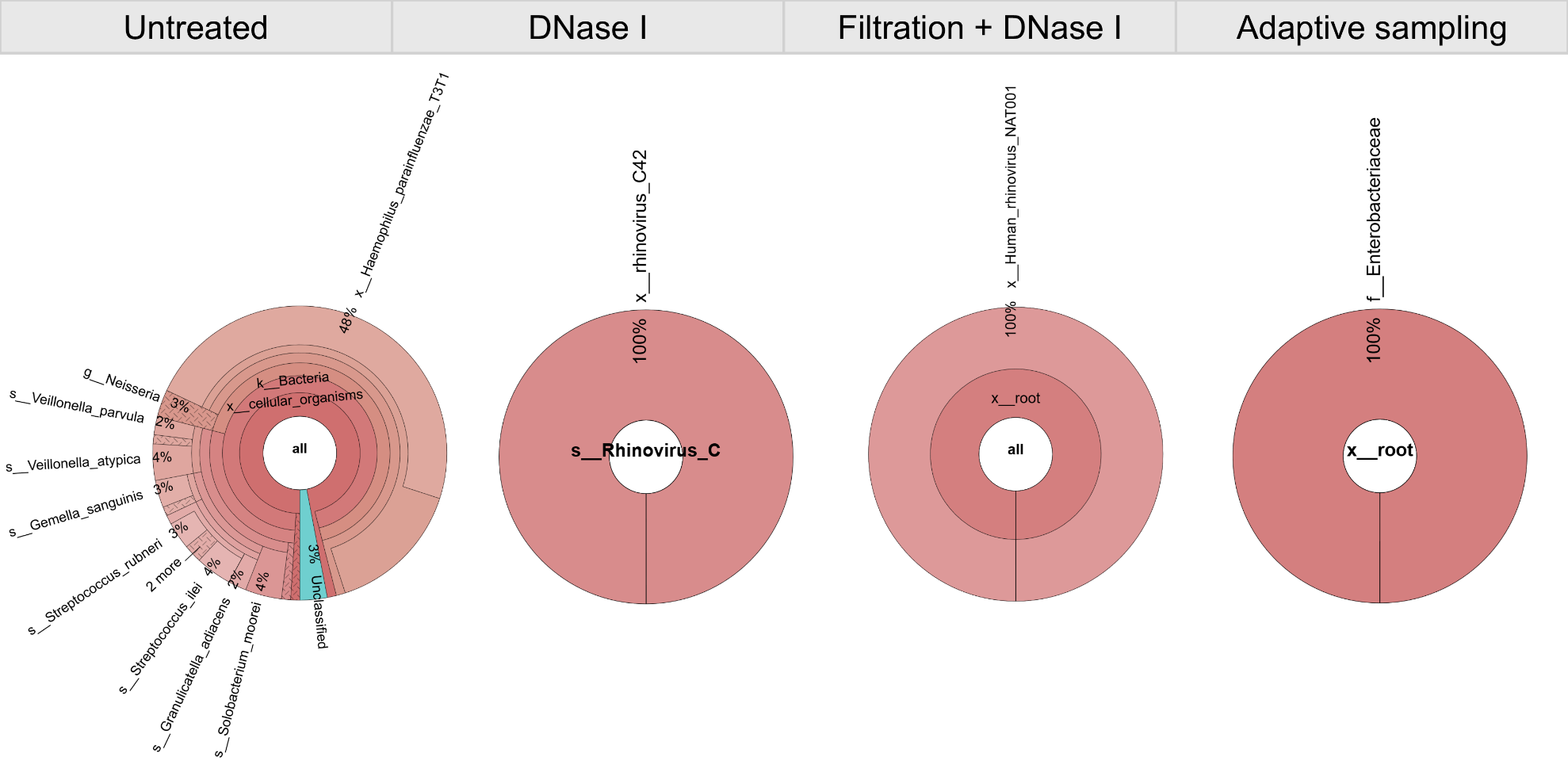

**Supplementary Figure S3.** Taxonomic composition of sample S4 following four host depletion strategies visualized using Krona plots.

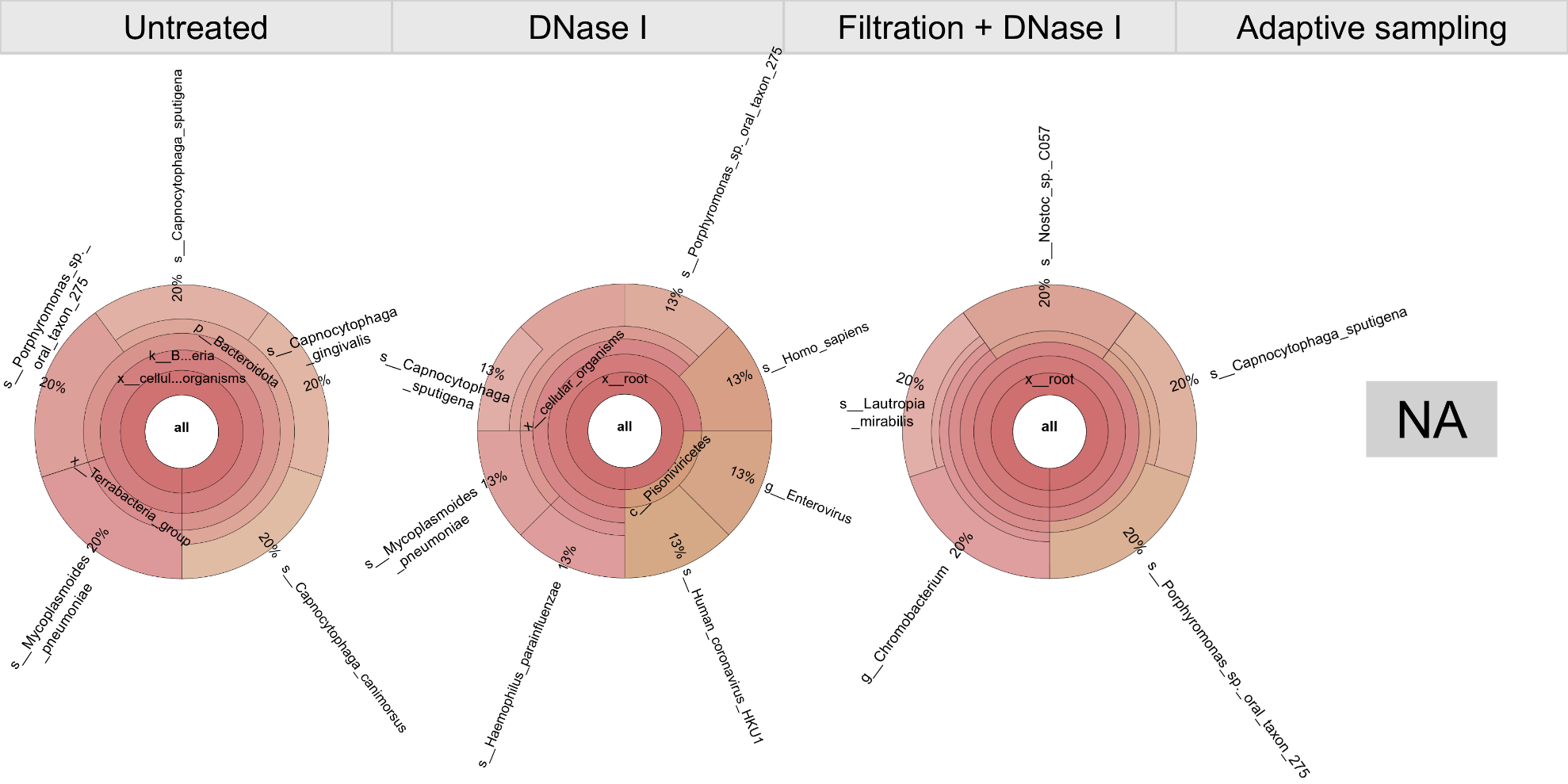

**Supplementary Figure S4.** Taxonomic composition of sample S5 following four host depletion strategies visualized using Krona plots. NA: not available.

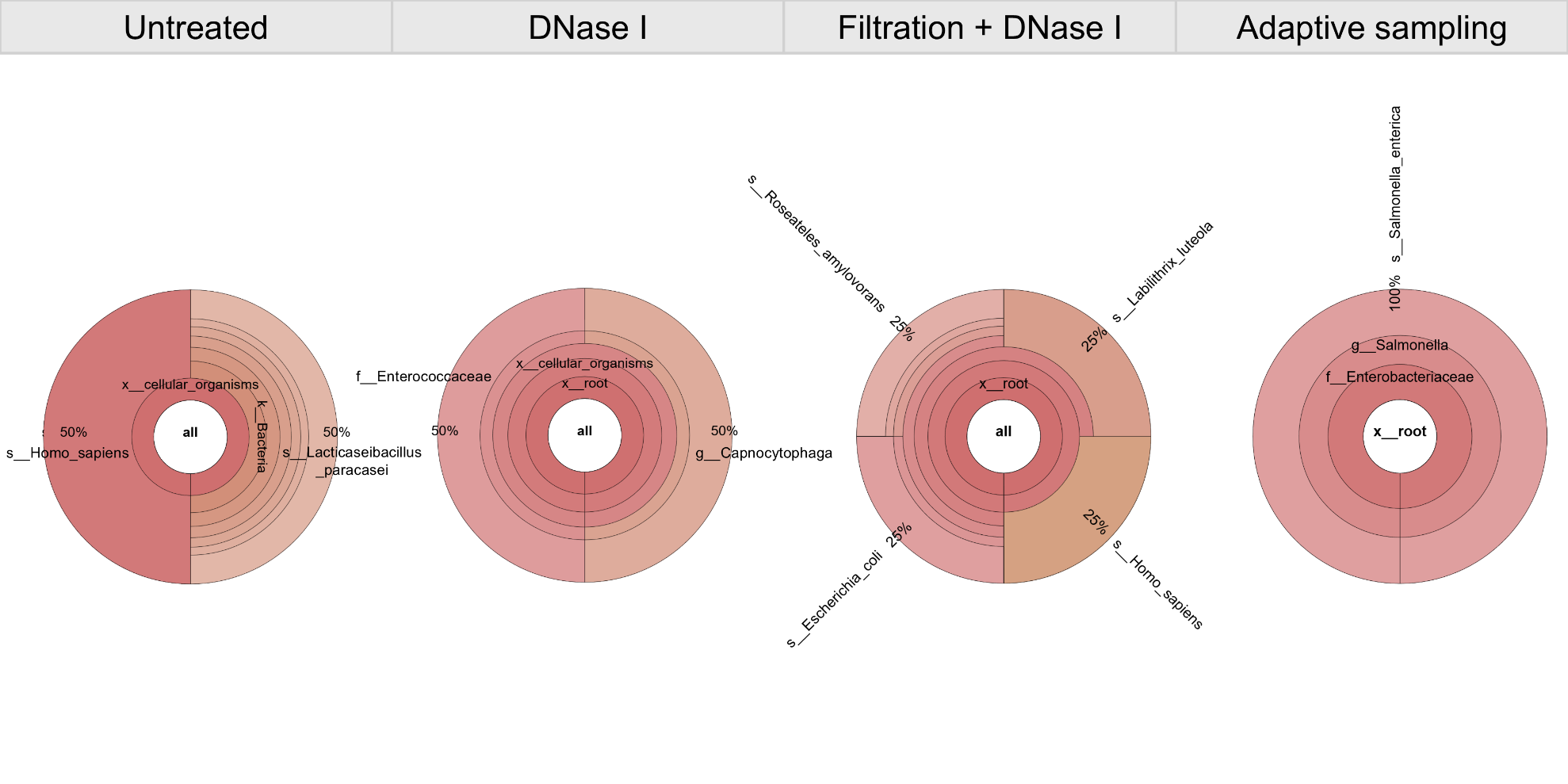

**Supplementary Figure S5.** Taxonomic composition of sample S6 following four host depletion strategies visualized using Krona plots.
